# ROS are evolutionary conserved cell-to-cell signals

**DOI:** 10.1101/2022.08.19.504606

**Authors:** Yosef Fichman, Linda Rowland, Melvin J. Oliver, Ron Mittler

## Abstract

Cell-to-cell communication is fundamental to multicellular organisms and unicellular organisms living in a microbiome. A unique cell-to-cell communication mechanism that uses reactive oxygen species (ROS) as a signal (termed the ‘ROS wave’) was recently identified in flowering plants. Here we report that a ROS-mediated cell-to-cell signaling process, like the ROS wave, can be found in ferns, mosses, unicellular algae, amoeba, mammalian cells, and isolated hearts. We further show that this process can be triggered by a localized stress treatment or H_2_O_2_ application and blocked by inhibition of NADPH oxidases, and that in unicellular algae, it communicates important stress-response signals between cells. Taken together, our findings suggest that cell-to-cell ROS signaling evolved before unicellular and multicellular organisms diverged. The finding of a ROS wave-like signaling process in mammalian cells further contributes to our understanding of different diseases and could impact the development of new drugs that target cancer or heart disease.

## Introduction

Cells communicate with each other in many ways. The exchange of signals between cells is thought to have evolved as part of a mechanism to control cell density (*e.g*., quorum sensing; QS)^1,2^, and/or as part of a response mechanism to changes in environmental conditions (*e.g*., in cells living in a biofilm)^3,4^. Cell-to-cell communication can occur between cells of the same species or organism (*e.g*., in a multicellular organism)^5,6^, as well as between cells of different organisms (*e.g*., during interactions of cells with different bacterial or fungal pathogens)^7,8^. Two of the most conserved cell-to-cell signaling pathways are paracrine, secretion of signals that are perceived by neighboring cells, and juxtacrine, communication between cells connected via tunneling nano tubes, gap junctions, or plasmodesmata^9–11^. Examples of these signaling pathways can be found in archaea, bacteria, unicellular eukaryotes, and multicellular organisms^12^. With the evolution of the vascular and nervous systems, paracrine signaling was extended to endocrine signaling, the release of signals into the bloodstream, and paracrine and juxtacrine signaling began to contribute to long distance signaling as part of electrical and chemical synapses^5,6^. In addition, autocrine signaling evolved as a mechanism for cells to regulate their own function^13^. Cell-to-cell signaling therefore plays an important role in coordinating the developmental and environmental responses of groups of neighboring cells, as well as cells that are remotely connected via blood vessels or neurons.

In vascular plants, that lack true nervous and circulatory systems, local cell-to-cell communication is primarily mediated by paracrine (via the apoplast and cell wall) and juxtacrine (via plasmodesmata; PD) signaling, while long distance signaling is mediated through the vascular system (phloem and xylem vessels, as well as their associated cells)^14,15^. Among the key signals exchanged by plant cells are different metabolites, small RNA and peptide molecules, hormones, calcium, volatile compounds, hydraulic pressure signals, membrane potential changes (electric signals), and reactive oxygen species (ROS)^14–18^. A unique cell-to-cell signaling mechanism discovered in plants is the ‘ROS wave’^19–21^. The ROS wave is an auto-propagating cell-to-cell signaling pathway in which the initiating cell activates extracellular (apoplastic) ROS production via respiratory burst oxidase homologs [RBOHs; the plant homologs of NADPH oxidases (NOXs) of mammalian cells]^19–21^. The ROS produced by this cell is then sensed by neighboring cells and triggers in them a similar enhanced ROS production process, resulting in the activation of a cascade or wave of cell-to-cell ‘enhanced ROS production state’ that spreads from the initiating cells/tissues over long distances, sometimes spanning the entire length of the plant^20,21^. The ROS wave was also recently shown to spread from one plant to another, when plants physically touch each other and conditions are sufficiently humid^22^. In addition to plasma membrane-localized RBOHs, the ROS wave requires the function of PD-localized proteins, different calcium channels, calcium signaling proteins, phytochrome B, and the plasma membrane (PM)-localized apoplastic ROS receptor H_2_O_2_-induced Ca^2+^ increases 1 (HPCA1) protein^21,23,24^. Numerous studies have shown that the ROS wave is linked to cell-to-cell calcium and electric signaling and that it is required for plant acclimation to stress (*e.g*.,^17,21,24–27^).

Because ROS are thought to have been present on Earth for billions of years and to have accompanied the evolution of unicellular and multicellular organisms^28–32^, we hypothesized that ROS, and perhaps a ROS-driven cell-to-cell signaling mechanism, like the ROS wave, could have evolved early during the evolution of life on Earth. To test this hypothesis, we studied whether ROS, and in particular an active auto-propagating process of cell-to-cell signaling mediated by ROS (like the ROS wave in flowering plants), can be found in plant lineages that diverged early in the plant phylogeny, including gymnosperms, ferns, lycophytes, bryophytes, multi and unicellular algae, as well as across the animal lineages in amoeba and mammalian cells. Here we show that a ROS-mediated cell-to-cell signaling process can be found in many of these organisms and cells, suggesting that ROS are evolutionarily conserved cell-to-cell signals.

## Results

### ROS as rapid systemic signals in vascular plants

We previously identified and studied the ROS wave in several different angiosperm species including Arabidopsis, rice, tomato, maize, dandelion, and wheat^20,22,24,33^. However, whether ROS can function as a rapid systemic signal in gymnosperms, as well as other plant lineages is unknown. Here we show that a rapid systemic ROS signal can be triggered by a local treatment of high light (HL) stress in *Pinus sylvestris* (Pine, a Gymnosperm), *Azolla filiculoides* (Azolla, a fern), and *Selaginella moellendorffii* (Selaginella, a Lycopodiaceae; Figure 1). The average rate of the ROS signal measured in these organisms is 0.14 cm min^-1^ (Table S1), faster than the diffusion rate of H_2_O_2_ in water or biological systems (the diffusion coefficient of H_2_O_2_ in water or tissues is in the range of 1-2.5 10^-5^ cm^2^ min^-1^)^34–36^, supporting the active nature of this systemic ROS signal. The findings presented in Figure 1 suggest that rapid systemic ROS signaling is conserved across several different members of the vascular plant family and may have evolved before these lineages diverged from a common ancestor.

**Figure 1.**
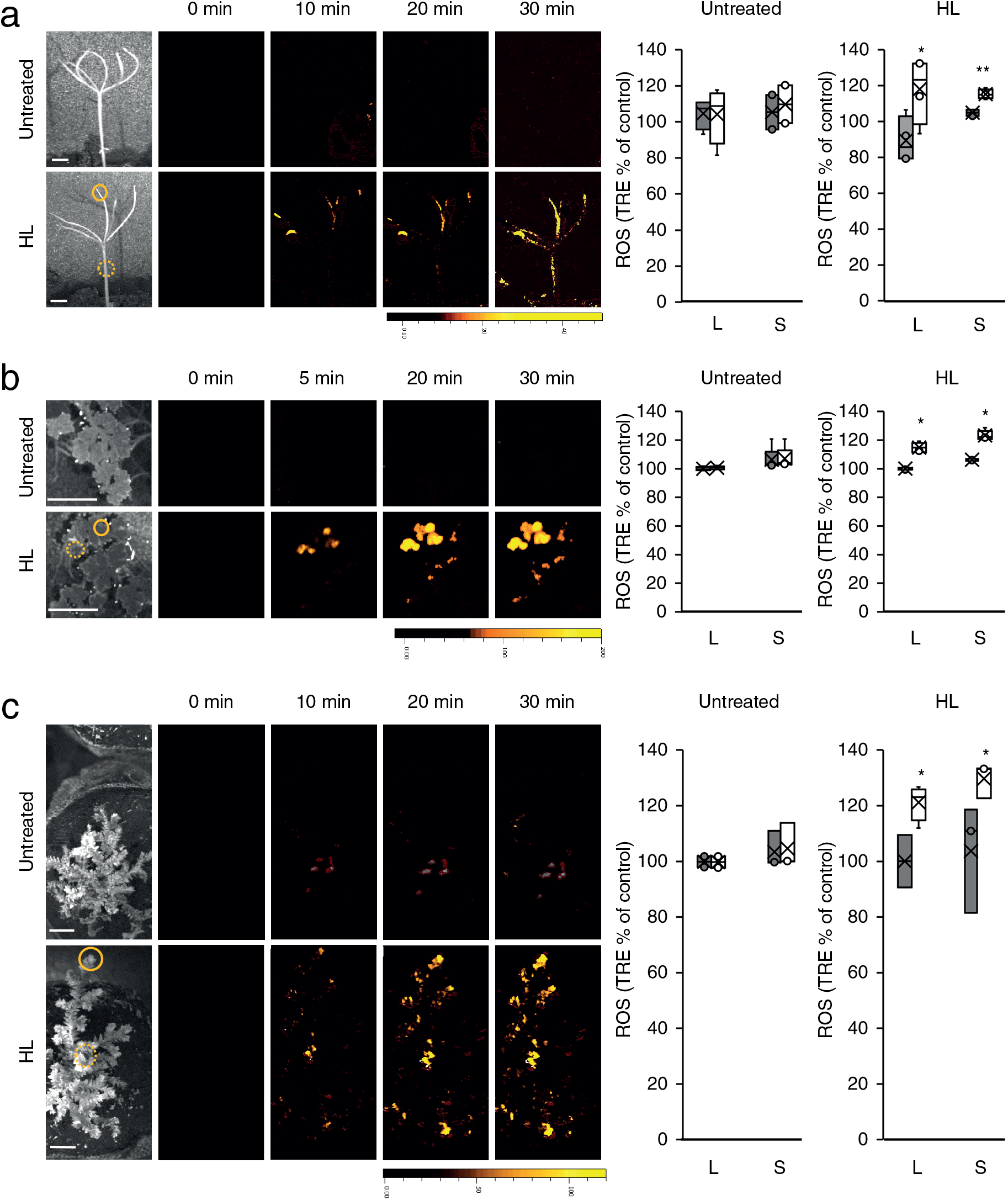
ROS as rapid systemic signals in *Pinus sylvestris, Azolla filiculoides*, and *Selaginella moellendorffii*. **(a)** *Pinus sylvestris* plants were subjected to a high light (HL) stress treatment applied to a single leaf, and ROS accumulation was imaged, using H2DCFDA, in whole plants (local and systemic tissues, indicated with solid and dashed circles, respectively). Representative time-lapse images of whole plant ROS accumulation in treated and untreated *Pinus sylvestris* plants are shown alongside bar graphs of combined data from all plants used for the analysis at the 0- and 30-min time points (local and systemic). **(b)** Same as a but for *Azolla filiculoides*. **(c)** Same as a, but for *Selaginella moellendorffii*. All experiments were repeated at least 3 times with 10 plants per experiment. Data is presented as box plot graphs; X is mean ± S.E., N=30, *P < 0.05, **P < 0.01, Student t-test. Scale bar, 1 cm. Abbreviations: H_2_DCFDA, 2’,7’-dichlorodihydrofluorescein diacetate; HL, high light; L, local; ROS, reactive oxygen species; S, systemic; TRE, total radiant efficiency.

### Rapid systemic ROS signaling in non-vascular plants

Although the ROS wave was shown to primarily propagate through the vascular bundles of Arabidopsis in response to a local HL stress treatment^37^, in response to a local wounding or heat stress treatments it could also propagate through non-vascular tissues (*i.e*., mesophyll cells)^38^. We therefore extended our study to include Bryophytes (hornworts, liverworts, and mosses) that emerged from an early split in the diversification of land plants and lack lignified vascular tissues^39^. As shown in Figure 2, a localized treatment of HL caused the activation of a rapid systemic ROS signal in *Anthoceros agrestis* (a Hornwort), *Marchantia polymorpha* (a Liverwort), and *Physcomitrium patens* (Physcomitrella, a Moss), with an average velocity of 0.13 cm min^-1^ (Table S1). These findings suggest that rapid systemic ROS signals are conserved across all land plants and can spread in a systemic manner within or between plants, even in plants that lack a true vascular tissue.

**Figure 2.**
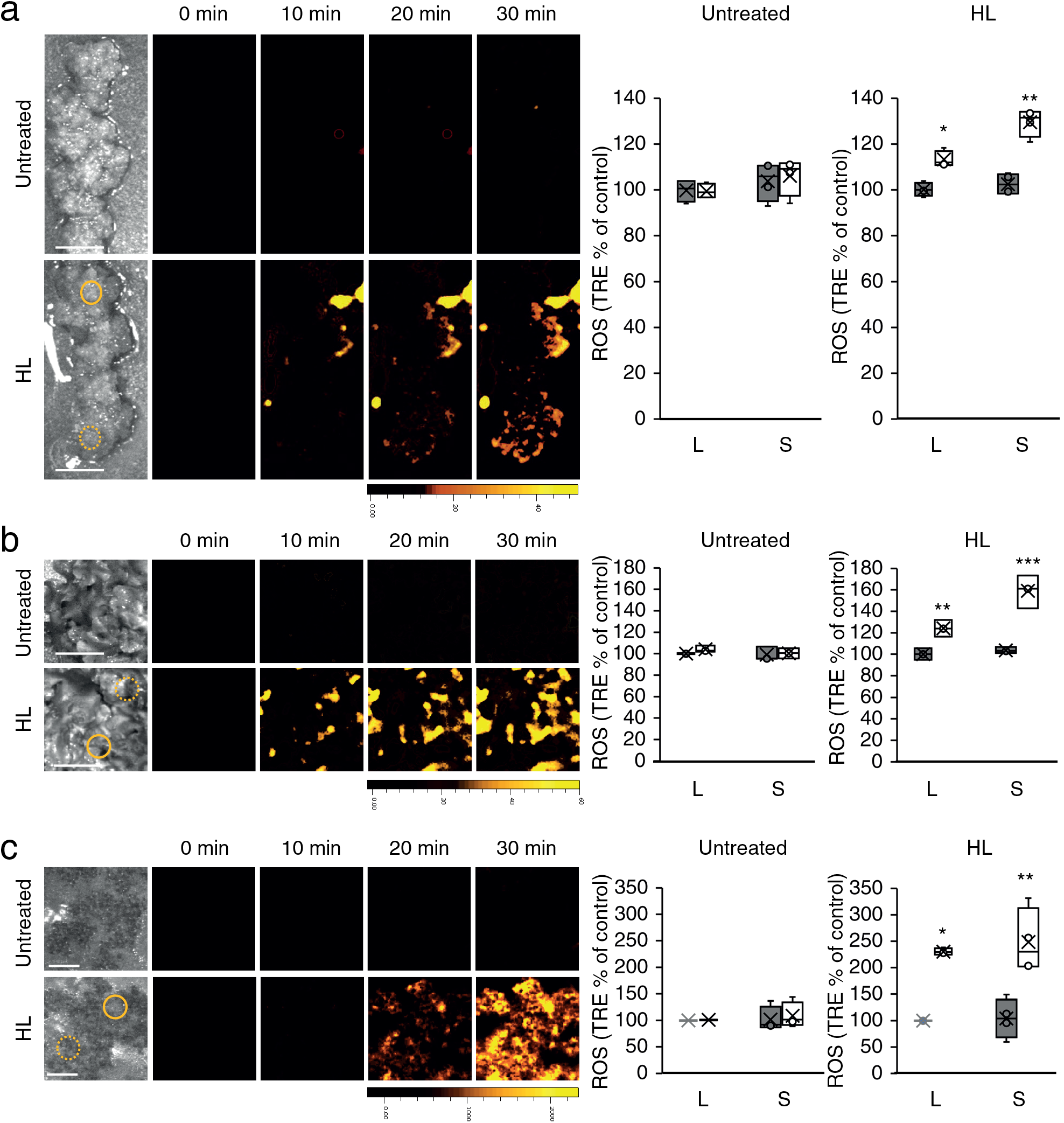
Rapid systemic ROS signaling in non-vascular plants. **(a)** *Anthoceros agrestis* plants were subjected to a high light (HL) stress treatment applied to a single leaf and ROS accumulation was imaged using H2DCFDA, in whole plants (local and systemic tissues, indicated with solid and dashed circles, respectively). Representative time-lapse images of whole plant ROS accumulation in treated and untreated *Anthoceros agrestis* plants are shown alongside bar graphs of combined data from all plants used for the analysis at the 0- and 30-min time points (local and systemic). **(b)** Same as a but for *Marchantia polymorpha*. **(c)** Same as a, but for *Physcomitrium patens*. All experiments were repeated at least 3 times with 10 plants per experiment. Data is presented as box plot graphs; X is mean ± S.E., N=30, *P < 0.05, **P < 0.01, ***P < 0.001, Student t-test. Scale bar, 1 cm. Abbreviations: H2DCFDA, 2’,7’-dichlorodihydrofluorescein diacetate; HL, high light; L, local; ROS, reactive oxygen species; S, systemic; TRE, total radiant efficiency.

### Rapid systemic ROS signaling in multicellular algae and unicellular organisms

The findings that rapid systemic ROS signals are conserved among many land plants (Figures 1 and 2)^20,22,24,33^ prompted us to test whether they can also be found in multicellular (*Chara vulgaris)* and unicellular (*Chlamydomonas reinhardtii*) algae. Multicellular and unicellular algae are known to use extracellular ROS as signals and contain NOX/RBOH enzymes^40,41^. As shown in Figures 3a and 3b, a localized treatment of HL stress caused the activation of a rapid systemic ROS response in *Chara vulgaris* grown in liquid medium (velocity of 0.14 cm min^-1^; Table S1), as well as in *Chlamydomonas reinhardtii* grown as a lawn on an agar plate (velocity of 0.12 cm min^-1^; Table S1). These findings suggest that ROS can be used as rapid systemic signals even by unicellular organisms and may have evolved as a cell-to-cell signaling mechanism before multicellularity emerged.

**Figure 3.**
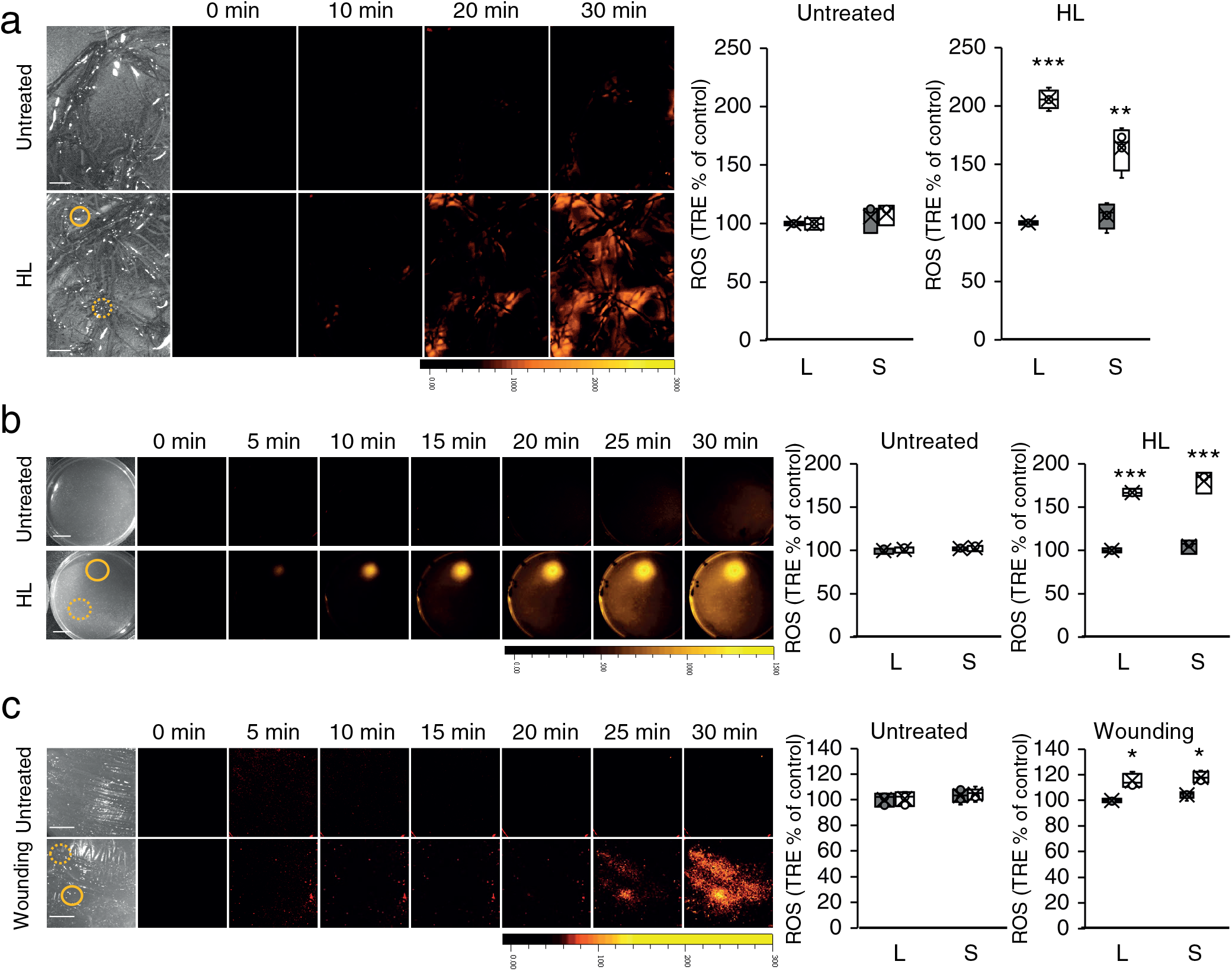
Rapid systemic ROS signaling in multicellular algae and unicellular organisms. **(a)** *Chara vulgaris*, a multicellular alga, was subjected to a high light (HL) stress treatment applied to a single branch, and ROS accumulation was imaged, using H2DCFDA, in several branches of the alga (local and systemic tissues, indicated with solid and dashed circles, respectively). Representative time-lapse images of whole alga ROS accumulation in treated and untreated *Chara vulgaris* are shown alongside bar graphs of combined data from all plants used for the analysis at the 0- and 30-min time points (local and systemic). **(b)** An agar plate containing a lawn of *Chlamydomonas reinhardtii*, a unicellular alga, was treated with a focused beam of HL (local treatment), and ROS accumulation was imaged, using H2DCFDA, in the entire plate (local and systemic tissues are indicated with solid and dashed circles, respectively). Representative time-lapse images of whole plate ROS accumulation of treated and untreated *Chlamydomonas* lawn are shown alongside bar graphs of combined data from all plants used for the analysis at the 0- and 30-min time points (local and systemic). **(c)** Same as in b, but for an agar plate that contains a lawn of *Dictyostelium discoideum* that was treated locally with a heated rod (local). All experiments were repeated at least 3 times with 10 multicellular alga, or agar plates per experiment. Data is presented as box plot graphs; X is mean ± S.E., N=30, *P < 0.05, **P < 0.01, ***P < 0.001, Student t-test. Scale bar, 1 cm. Abbreviations: H2DCFDA, 2’,7’-dichlorodihydrofluorescein diacetate; HL, high light; L, local; ROS, reactive oxygen species; S, systemic; TRE, total radiant efficiency.

The findings that rapid systemic ROS signaling can be found in unicellular alga (Figure 3b) prompted us to test whether it can also be found in other unicellular organisms, such as the soil dwelling amoeba *Dictyostelium discoideum*, that contains NOX and uses extracellular ROS for signaling^41–43^. As shown in Figure 3c, a localized treatment of heat injury caused the activation of a rapid systemic ROS signal in the amoeba *Dictyostelium discoideum* grown on agar plates (velocity of 0.2 cm min^-1^; Table S1).

### Initiation and inhibition of the ROS wave in *Physcomitrella, Chlamydomonas*, and *Dictyostelium*

Two important characteristics of the ROS wave are that it can be triggered by a localized application of H_2_O_2_ and that once activated it can be blocked by the application of diphenyleneiodonium chloride (DPI; a broad-spectrum NOX/RBOH inhibitor) at a distant location away from its initiation site^19–21^. As shown in Figure 4, application of a 1 μl drop of 10 mM H_2_O_2_ to *Physcomitrella, Chlamydomonas*, or *Dictyostelium*, grown on agar plates, triggered a rapid systemic ROS signal. In contrast, application of mock buffer to cells, or application of 1 μl drop of 10 mM H_2_O_2_ to agar plates that contain no cells, did not result in a ROS signal. To further test for H_2_O_2_ diffusion on plates that had no cells grown on them (live cells are required for H2DCFDA to generate a signal)^44^, we used 2’,7’-dichlorodihydrofluorescein (DCF; OxyBurst; 50 μM)^19^ to image H_2_O_2_ in empty plates following the application of 1 μl of 10 mM, or 1 M H_2_O_2_. As shown in Figure S1, we could not detect the 1 μl of 10 mM H_2_O_2_, a concentration that resulted in a detectable ROS signal in plates with the different organisms (Figure 4) but were able to detect the 1 μl of 1 M H_2_O_2_. This finding suggests that the living cells of the different organisms shown in Figure 4 amplified the 1 μl of 1 mM H_2_O_2_ to detectable levels.

**Figure 4.**
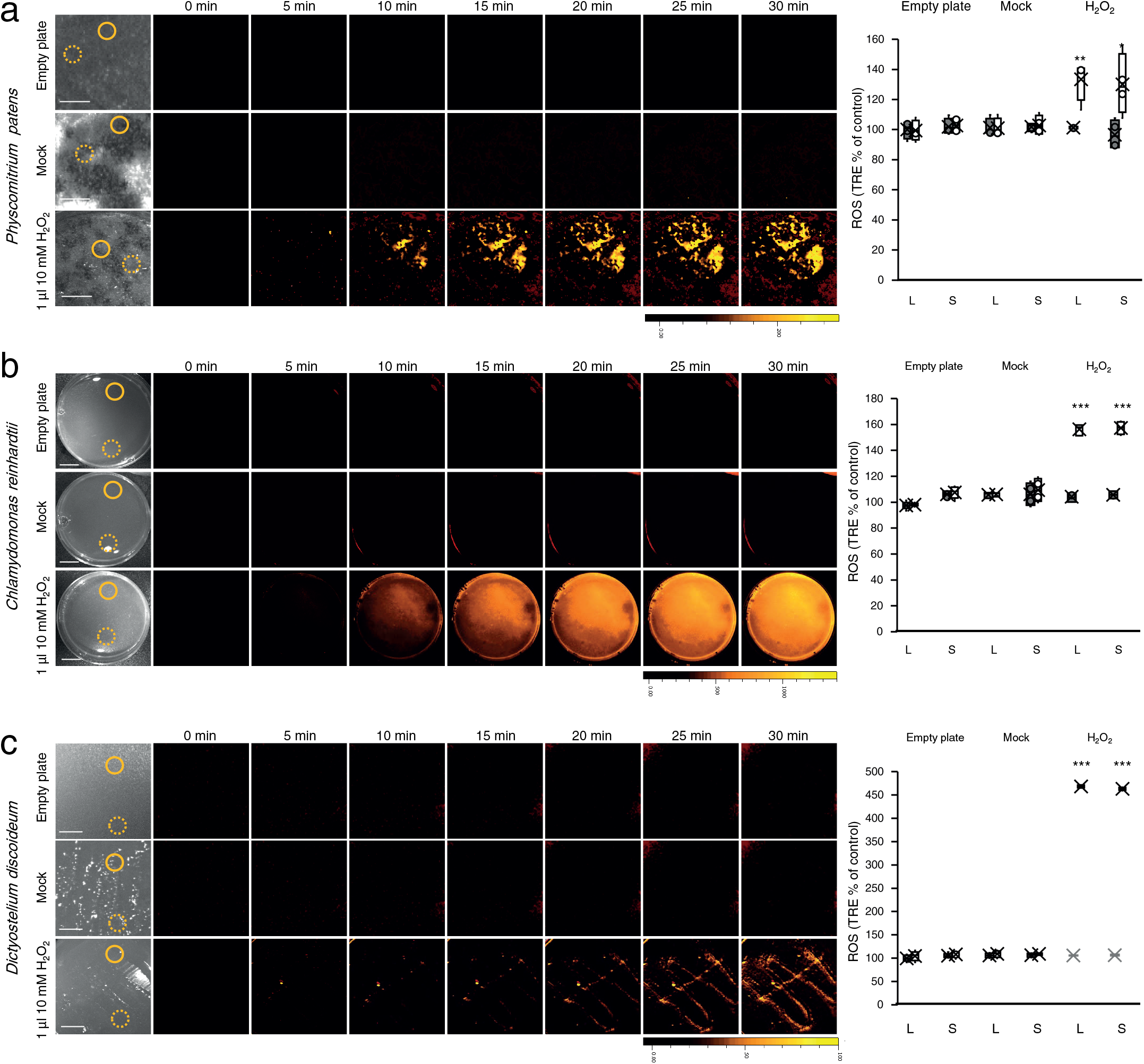
Activation of the rapid systemic ROS signal in *Physcomitrium patens, Chlamydomonas reinhardtii*, and *Dictyostelium discoideum* by a local application of H_2_O_2_. **(a)** An agar plate without *Physcomitrium patens* (top), and agar plates with a lawn of *Physcomitrium patens* (middle and bottom), were treated with 1 μl of 1 mM H_2_O_2_ (top and bottom), or 1 μl of media (mock, middle), and ROS accumulation was imaged using H2DCFDA in the entire plate. Representative time-lapse images of whole plate ROS accumulation are shown alongside bar graphs of combined data from all plates used for the analysis at the 0- and 30-min time points (local and systemic). Local and systemic sampling locations are indicated with solid and dashed circles, respectively. **(b)** Same as a, but for plates with or without a lawn of *Chlamydomonas reinhardtii*. **(c)** Same as in a, but for plates with or without *Dictyostelium discoideum*. All experiments were repeated at least 3 times with 10 agar plates per experiment. Data is presented as box plot graphs; X is mean ± S.E., N=30, *P < 0.05, **P < 0.01, ***P < 0.001, Student t-test. Scale bar, 1 cm. Abbreviations: H2DCFDA, 2’,7’-dichlorodihydrofluorescein diacetate; HL, high light; L, local; ROS, reactive oxygen species; S, systemic; TRE, total radiant efficiency.

To test the effect of the RBOH/NOX inhibitor DPI on the propagation of the rapid systemic ROS signal in *Physcomitrella, Chlamydomonas*, or *Dictyostelium*, grown on agar plates, we applied water or DPI (50 μM) in a 1 mm wide strip of agar to the middle of plates, waited 5 min and applied HL or heat injury at one end of the plate (Figure 5a). As shown in Figure 5b, application of DPI (but not water) to the middle of the plate blocked the progression of the cell-to-cell ROS signal. The findings presented in Figures 4 and 5 suggest that the rapid systemic ROS signal, observed as propagating from the initial site of HL or heat injury on plates in the different organisms shown in Figures 2–4, is auto-propagating and resembles the ROS wave discovered and studied in *Arabidopsis thaliana*^19–21,37,38^.

**Figure 5.**
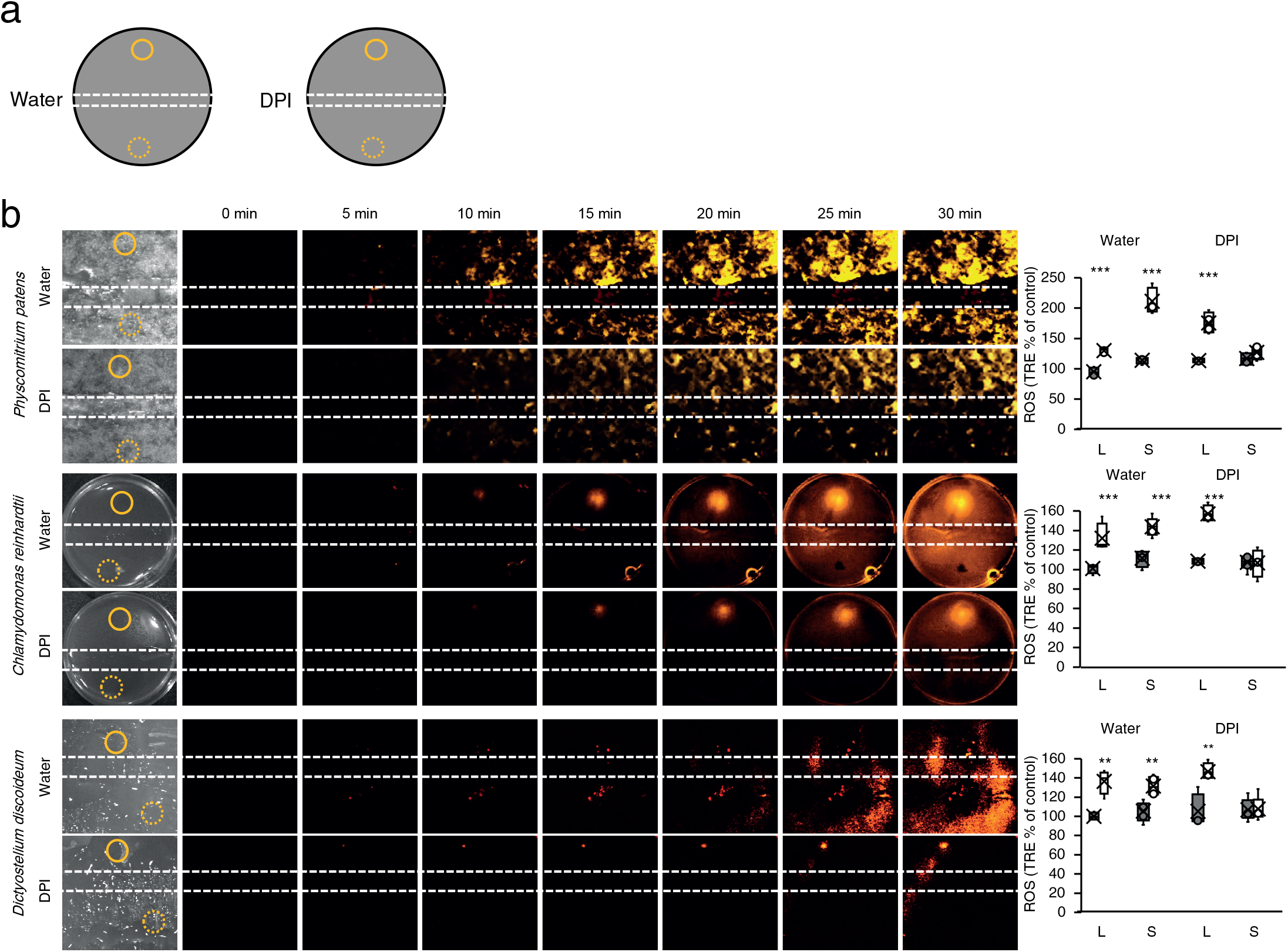
Inhibition of the rapid systemic ROS signal in *Physcomitrium patens, Chlamydomonas reinhardtii*, and *Dictyostelium discoideum* by DPI. **(a)** A diagram showing the experimental design used. A strip of agar containing water or the NOX/RBOH inhibitor DPI (50 μM) was layered over the middle of agar plates containing a lawn of the different organisms growing on them. Following a 5 min incubation one side of the plate was treated with a focused beam of high light (HL) or touched with a heated rod (local) and ROS accumulation was imaged using H2DCFDA in the entire plate. Local and systemic locations used for signal quantification are indicated with solid and dashed circles, respectively. **(b)** Representative time-lapse images of whole plate ROS accumulation are shown alongside bar graphs of combined data from all plates used for the analysis at the 0- and 30-min time points (Water or DPI treated *Physcomitrium patens*, *Chlamydomonas reinhardtii*, or *Dictyostelium discoideum*; top, middle, and bottom, respectively; local and systemic). All experiments were repeated at least 3 times with 10 agar plates per experiment. Data is presented as box plot graphs; X is mean ± S.E., N=30, **P < 0.01, ***P < 0.001, Student t-test. Scale bar, 1 cm. Abbreviations: DPI, diphenyleneiodonium; H2DCFDA, 2’,7’-dichlorodihydrofluorescein diacetate; HL, high light; L, local; NOX, NADPH oxidase; RBOH, respiratory burst oxidase homolog; ROS, reactive oxygen species; S, systemic; TRE, total radiant efficiency.

### Functional analysis of the rapid cell-to-cell systemic ROS signal in *Chlamydomonas reinhardtii*

To test whether the systemic ROS signal observed in the unicellular alga *Chlamydomonas reinhardtii* (Figures 3–5) can transfer a biologically relevant signal between different cells that are at two ends of the signal path (*i.e*., local and systemic, on two ends of the agar plate; Figure 6a), we applied HL stress to one corner of a lawn of *Chlamydomonas* cells and sampled these cells, as well as cells that are on the other end of the plate (*i.e*., not subjected to the HL stress treatment; Figure 6a). We then tested the expression of a HL-response small heat shock protein (sHSP17.1) in cells located at the two ends of the signal path following the application of HL stress to one end of the plate (stressed cells; local). As shown in Figure 6b, the expression of sHSP17.1 was elevated at both ends of the signal path (local and systemic; Figure 6a) suggesting that the ROS wave signal connecting these cells is conveying a stress response signal from the stressed cells (local) to the non-stressed cells (systemic) that did not sense the stress.

**Figure 6.**
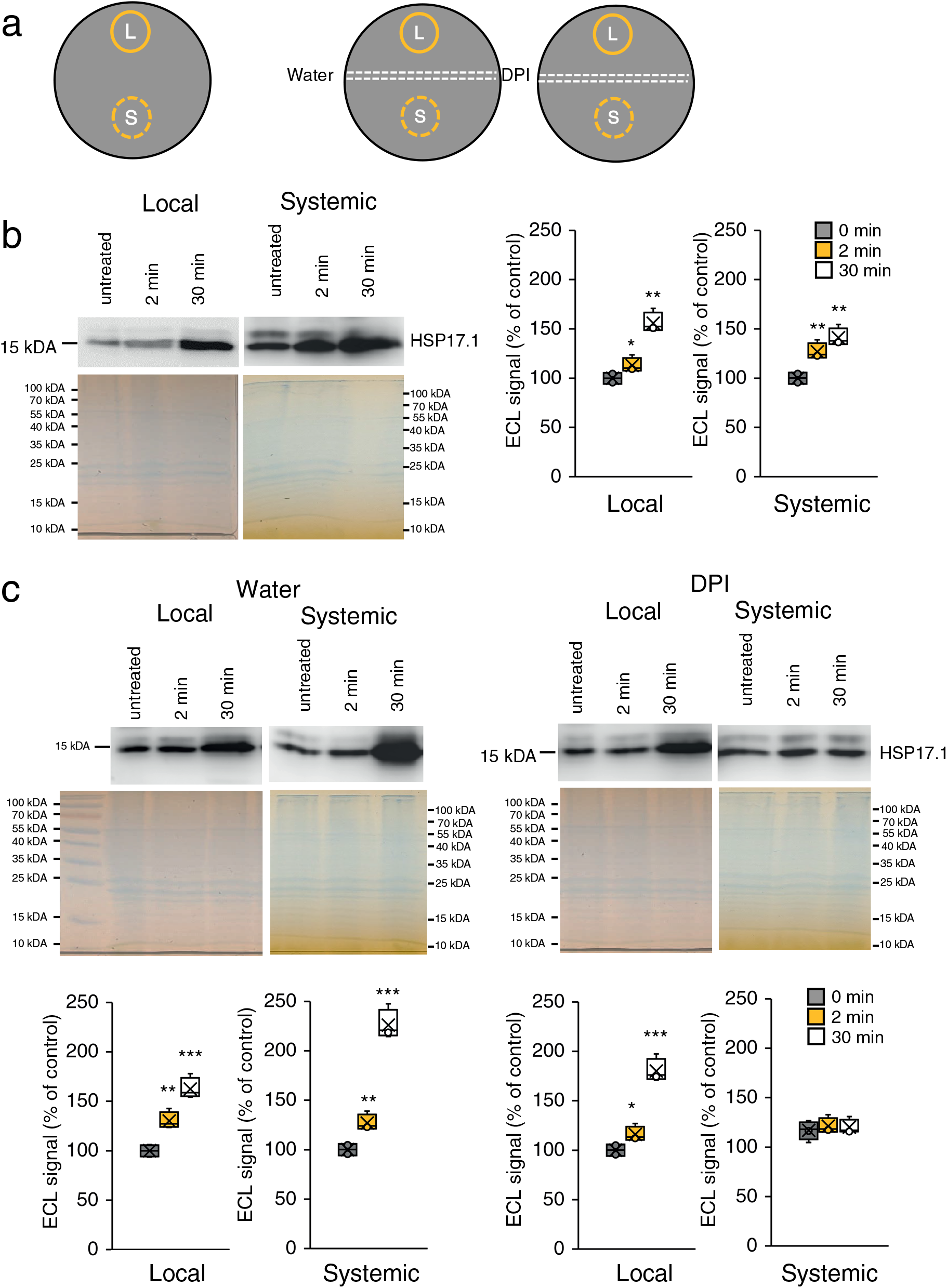
Functional analysis of the cell-to-cell rapid ROS signal in *Chlamydomonas reinhardtii*. **(a)** Diagrams showing the experimental designs used. Left: A lawn of *Chlamydomonas reinhardtii* growing on an agar plate was subjected to a focused beam of high light (HL) stress and the area subjected to the beam (local, L), as well as an area that is on the other end of the plate (systemic, S) were sampled at different times following the application of HL for analysis of the expression of the light stress response small heat shock protein 17.1 (sHSP17.1; This design was used for the experiments shown in B). Right: A strip of agar containing water or the NOX/RBOH inhibitor DPI (50 μM) was layered over the middle of agar plates containing the lawn of *Chlamydomonas reinhardtii* 5 min prior to the application of light stress and sampling (This design was used for the experiments shown in C). Local and systemic sampling locations are indicated with solid and dashed circles, respectively. **(b)** Representative protein blots and Coomassie-stained gels are shown alongside bar graphs of combined data from all blots used for the analysis at the 0-, 2-, and 30-min time points, for the expression of sHSP17.1 in the L and S locations, following the light stress treatment. **(c)** Same as in b, but for experiments in which a strip of agar containing water (left) or DPI (right) was layered over the middle of agar plates 5 min prior to the application of HL stress. All experiments were repeated at least 3 times with 10 agar plates per experiment. Data is presented as box plot graphs; X is mean ± S.E., N=30, *P < 0.05, **P < 0.01, ***P < 0.001, Student t-test. Scale bar, 1 cm. Abbreviations: DPI, diphenyleneiodonium; H2DCFDA, 2’,7’-dichlorodihydrofluorescein diacetate; HL, high light; L, local; NOX, NADPH oxidase; RBOH, respiratory burst oxidase homolog; ROS, reactive oxygen species; S, systemic; sHSP, small heat shock protein 17.1.

To test whether inhibiting the ROS wave in *Chlamydomonas* growing on an agar plate (Figure 5) would inhibit the transfer of the stress-response signal from one side of the plate to the other (Figure 6b), we applied DPI (50 μM) or water at the middle point between the stressed cells and the cells that did not sense the stress (similar to Figure 5; Figure 6a). As shown in Figure 6c, DPI, but not water, inhibited the expression of sHSP17.1 in the non-stressed (systemic) cells following the application of HL stress to the stressed cells (local) that are on the other end of the plate. In contrast, DPI or water did not inhibit the expression of sHSP17.1 at the stressed cells (local). These findings suggest that the cell-to-cell ROS signal connecting the two ends of the *Chlamydomonas* lawn growing on the agar plate transfers biologically relevant signals between cells.

### Cell-to-cell ROS signaling in mammalian cells

The findings that the ROS wave could be found in the amoeba *Dictyostelium discoideum* (Figure 3c), demonstrated that organisms that are outside the plant and algal lineages could use ROS as a cell-to-cell signaling mechanism. *Dictyostelium discoideum* was previously found to contain an ancestral form of NOX^28,41,43^, suggesting that other organisms that contain NOX enzymes (for example mammalian cells) could be using ROS for rapid cell-to-cell signaling. To test this possibility, we studied cell-to-cell ROS signaling in monolayers of human (*Homo sapiens)* epithelial breast cancer (MDA-MB-231), and rat (*Rattus norvegicus*) cardiomyocytes (H9c2), cells grown in culture. As shown in Figure 7a, wounding of a monolayer of MDA-MB-231 or H9c2 cells with a heated rod resulted in a ROS signal that spread from the injury site at a velocity of 0.28 or 0.2 cm min^-1^ (Table S1), respectively. In the case of H9c2 cells, the ROS signal appeared to travel away from the wound site and to do so in a directional manner, suggesting that the ROS signal observed is not simply ROS that are diffusing away from the initial wound site in all directions (Figure 7a; Movie S1). To determine that the spreading ROS signal was not a result of cell migration and/or monolayer detachment, we conducted microscopic analysis of H9c2 cells at the end the ROS imaging experiments. This analysis revealed that the wound site had an average diameter of 300-600 μm and that cells did not detach or migrate away from it (Figure S2). Moreover, migration of cells away from the wound site would have been much slower (1–10 μm min^-1^)^45^ than the rate of the signal observed (0.2 cm min^-1^; Table S1). In addition, to test that H_2_O_2_ can be detected in all H9c2 cells on the plates (not only the cells observed generating ROS after wounding; Figure 7a), we treated the entire plate with a final concentration of 1 mM H_2_O_2_ at the end of the experiment (Figure S3). This analysis revealed that all cells on the plate were able to uptake the H2DCFDA dye, and that H_2_O_2_ that was provided to the media and entered cells generated a signal [for H2DCFDA to generate a fluorescent signal at the rapid rate the signal is developing (Figure 7a), it has to enter live cells]^44^. To test whether the ROS signal observed in MDA-MB-231 or H9c2 is associated with NOX function, we applied DPI (50 μM) or buffer (mock) to these cells prior to the heat injury. As shown in Figure 7b, DPI, but not buffer, caused the complete suppression of the ROS signal.

**Figure 7.**
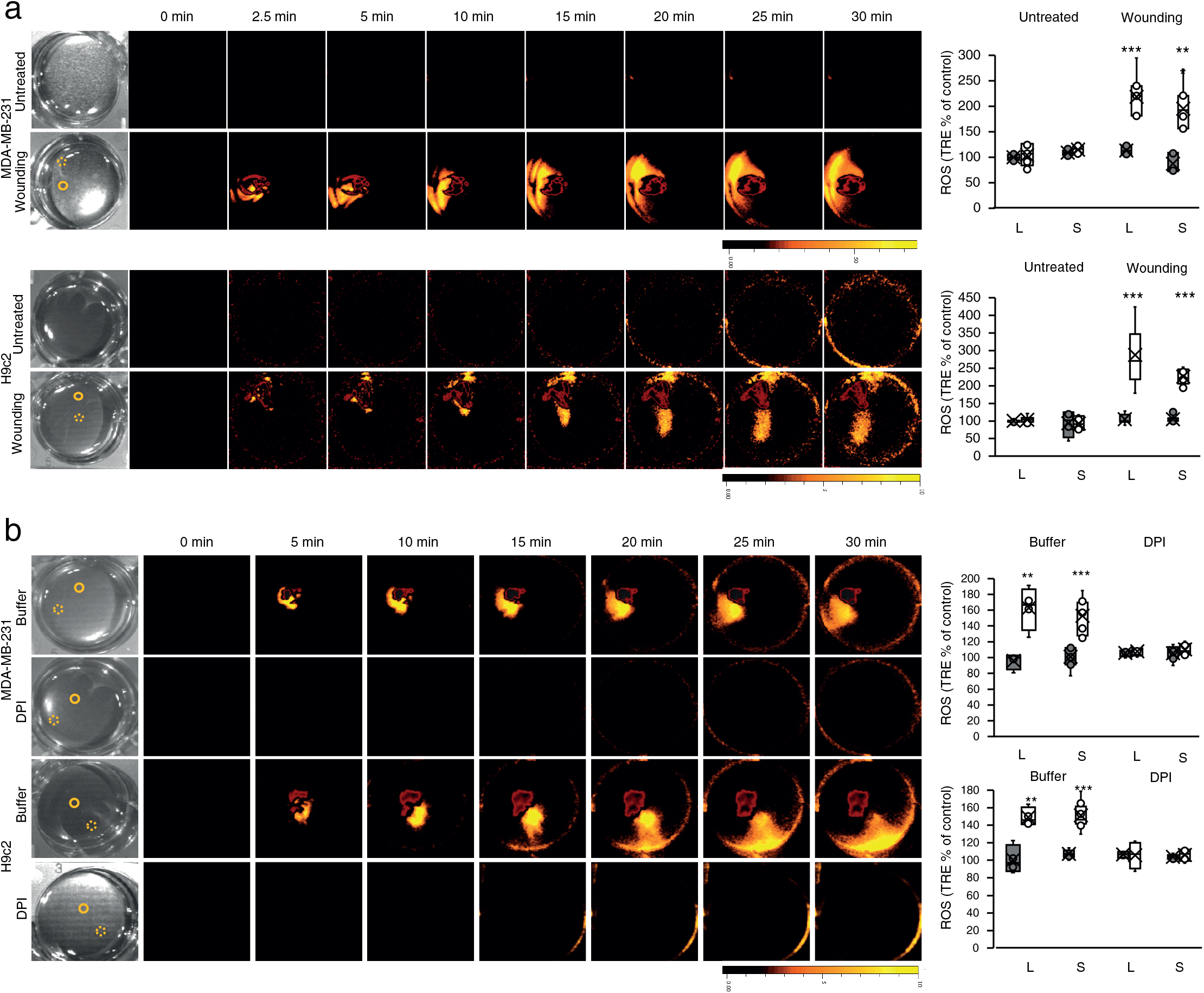
Rapid systemic ROS signaling in mammalian cells. **(a)** Monolayers of human epithelial breast cancer (MDA-MB-231; top), and rat cardiomyocytes (H9c2; bottom), cells grown in culture were treated with a heated metal rod to induce injury (average injury diameter is 120 μm) and ROS accumulation was imaged using H2DCFDA in the entire plate. Representative time-lapse images of whole plate ROS accumulation in treated and untreated plates are shown alongside bar graphs of combined data from all plates used for the analysis at the 0- and 30-min time points (local and systemic). Local and systemic sampling locations are indicated with solid and dashed circles, respectively. **(b)** Same as in a, except that buffer or DPI (50 μM) were added to the growth media of cells 10 min prior to the heat injury. All experiments were repeated at least 3 times with 10 agar plates per experiment. Data is presented as box plot graphs; X is mean ± S.E., N=30, **P < 0.01, ***P < 0.001, Student t-test. Scale bar, 1 cm. Abbreviations: DPI, diphenyleneiodonium; H2DCFDA, 2’,7’-dichlorodihydrofluorescein diacetate; L, local; ROS, reactive oxygen species; S, systemic; TRE, total radiant efficiency.

### Cell-to-cell ROS signaling in isolated mice hearts

The finding of a ROS wave-like signaling process in rat cardiomyocytes grown in culture (Figure 7) prompted us to test whether a similar process could be found in isolated hearts obtained from mice. For this purpose, we collected hearts from 9-10-weeks-old female and male C57BL/6J mice and incubated them with H_2_DCFDA for 2 hours. We then wounded the isolated hearts using an open-flamed-heated rod and imaged ROS responses in the entire isolated heart. As shown in Figures 8a, S4, and Movie S2, wounding of isolated hearts from female or male mice resulted in a ROS response that originated from the point of wounding and progressed at an average rate of 0.106 cm min^-1^ (Table S1) to other parts of the heart. As shown in Figures 8b and S4, this process was inhibited by DPI application to hearts 30 min prior to wounding. These findings suggest that a ROS wave-like signaling process could occur in isolated heart tissue.

**Figure 8.**
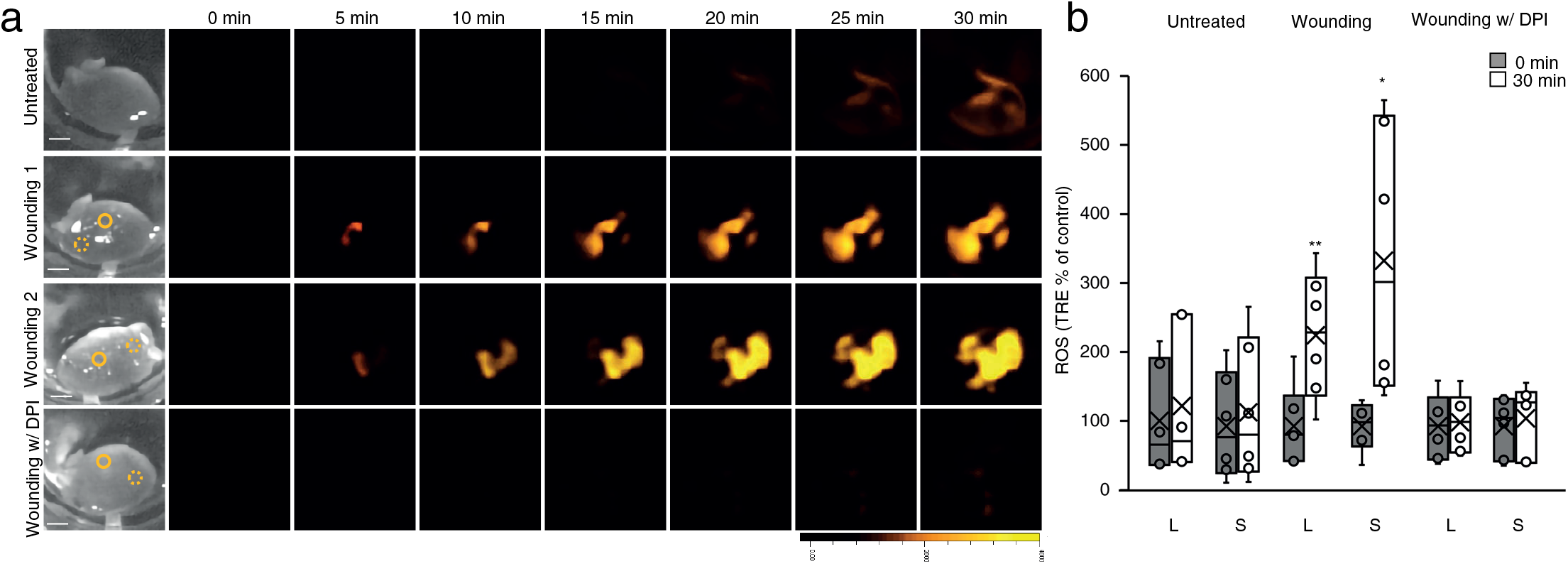
Wound-induced rapid systemic ROS signaling in isolated mice hearts. Mice hearts, surgically removed from 9-10-week-old male and female C57BL/6J mice, were untreated, or treated with a heated metal rod (to induce injury), in the presence or absence of DPI (50 μM), and ROS accumulation was imaged using H2DCFDA in the entire hearts. Representative time-lapse images of whole heart ROS accumulation **(a)** are shown alongside bar graphs of combined data from all hearts used for the analysis at the 0- and 30-min time points **(b)**. Local and systemic sampling locations are indicated with solid and dashed circles, respectively. All experiments were repeated at least 3 times with 3 female and 3 male mice per experiment. Data is presented as box plot graphs; X is mean ± S.E., N=30, **P < 0.01, ***P < 0.001, Student t-test. Scale bar, 1 cm. Female- and male-specific ROS responses are shown in Figure S4. Abbreviations: DPI, diphenyleneiodonium; H2DCFDA, 2’,7’-dichlorodihydrofluorescein diacetate; L, local; ROS, reactive oxygen species; S, systemic; TRE, total radiant efficiency; w/, with.

## Discussion

The steady-state level of ROS in cells can vary in response to different environmental conditions, the activation of developmental programs that cause active production of ROS, and/or a decline in the activity of ROS scavenging mechanisms^46–48^. ROS can also diffuse into or out of cells through aquaporins, further impacting intracellular ROS levels^47,48^. Because ROS affect the redox state of cells and can alter gene expression through oxidative post-translational modifications (oxi-PTMs) of different proteins^47^, they function as key signal transduction molecules involved in the sensing of stress and/or the activation of different developmental programs^48^.

Because ROS can be transported in or out of cells through aquaporins and/or other channels/pores, secretion of ROS by one cell can alter the redox state of a neighboring cell, thereby altering gene expression within it. This type of paracrine signaling is thought to play an important role in cell-to-cell signaling occurring in different communities of microorganisms, during algal blooms, during interactions between neurons, or during the recruitment of leukocytes to wound sites^49–53^. A central feature of this paracrine signaling mechanisms is however that it may depend on a gradient of ROS forming from the secreting cell(s) to the accepting cell(s). In addition, ROS can be scavenged in route from one cell to the other, limiting the range of this signaling pathway. Having the ability to amplify and maintain cell-to-cell ROS signals, like the auto-propagating ROS wave process in plants^19–21,37,38^, would improve the communication between cells in a biofilm/community/tissue and allow them to respond faster and in unison (Figure 9). Here we demonstrate that *Physcomitrella, Chlamydomonas*, and *Dictyostelium*, growing on agar plates, and mammalian cells growing in culture, indeed have the capacity to amplify and maintain cell-to-cell ROS signals (Figures 3–7; Table S1). We further show that cell-to-cell ROS signaling may also occur in isolated mice hearts (Figure 8). These findings suggest that the ROS wave signaling process, that was originally discovered in flowering plants^19–21,37,38^, could have originated early in evolution and is conserved between plants, microorganisms, and animals. In support of these data are the findings that the systemic ROS signal could be initiated by H_2_O_2_ (Figure 4), and that application of DPI at a location that is away from the initiation site could block it (Figure 5). Moreover, in H9c2 cells, the peak of the ROS signal propagated in a directional manner at a rate of 0.2 cm min^-1^ departing from its original initiation site and continuing to move away from it in a manner that clearly suggests it is not mediated by simple diffusion (Figures 7 and S3; Movie S1). Although the directionality of the ROS signal (Figures 7 and S3; Movie S1) could be related to cell polarity, it is unknown at present what is the mechanism that govern it. A similar directional spread of the ROS signal was also observed in isolated hearts (Figure 8; Movie S2), further supporting the notion that it is not a simple diffusion process. In addition, the rate of the ROS signal observed in hearts was also faster than that of H_2_O_2_ in biological systems or water (Table S1)^34–36^. Taken together, these findings suggest that the ROS signal observed in a monolayer of H9c2 cells (Figure 7), or isolated hearts (Figure 8), is driven by an active process of ROS production (that could be inhibited by DPI; Figures 7 and 8), involving multiple cells along its path (*i.e*., not only cells at and around the injury site).

**Figure 9.**
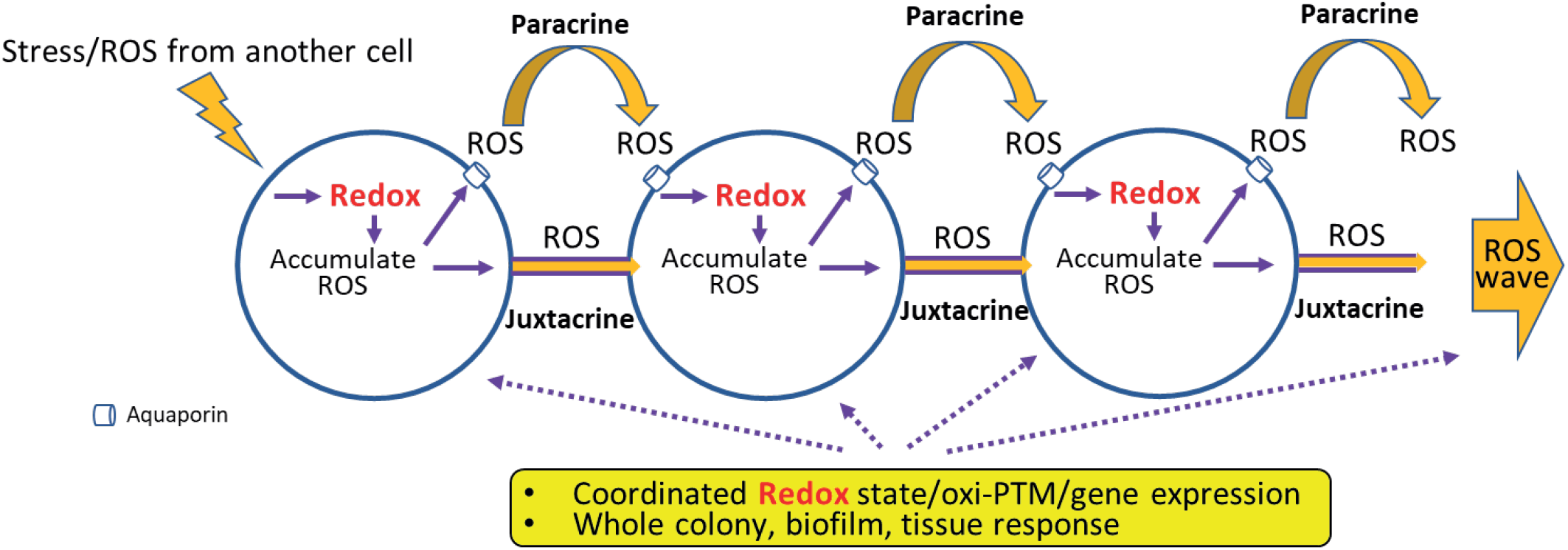
A model depicting the principle of the cell-to-cell ROS signaling pathway that is conserved between unicellular and multicellular organisms. An initiating cell that perceives a stress or ROS signal is shown on the left to undergo a change in redox levels that will cause the accumulation of ROS in that cell. This process is shown to cause the activation of a paracrine and juxtacrine ROS-mediated cell-to-cell signal transduction pathway. Each cell along the path of this pathway accumulates ROS maintaining the signal and transmitting it to the next. The resulting changes in redox in all cells involved in this signaling pathway activate gene expression via oxidative post translational modification (oxi-PTM) of different regulatory proteins, yielding a coordinated cellular response to the original stress or signal in the entire colony, biofilm, and/or tissue. Please see text for more details. Abbreviations: oxi-PTM, oxidative post translational modification; ROS, reactive oxygen species.

An interesting analogy to the cell-to-cell ROS wave signaling process of plants is the synchronized process of mitochondrial ROS flushes that occurs in mitochondria that are arranged in a lattice formation in muscle tissue (*e.g*.,^54,55^). In this process the release of ROS through pores from one mitochondrion results in the release of ROS from a neighboring mitochondrion spreading from mitochondria-to-mitochondria in a ROS-induced-ROS-release fashion^54,55^. In the context of our findings, that a similar process of cell-to-cell ROS signaling occurs in many different unicellular and multicellular organisms (Figures 1–8), it is possible that even ancient prokaryotic organisms that were the progenitors of mitochondria used this type of cell-to-cell signaling to orchestrate responses within a biofilm or a colony. It is also possible that mitochondria are actively involved in the cell-to-cell ROS signal found in H9c2 cells or hearts (Figures 7 and 8), and that NOXs such as NOX4 that is associated with mitochondria in mammalian cells^56^ are mediating this process. Further studies are needed to address the potential role of mitochondria in cell-to-cell ROS signaling.

In addition to paracrine signaling, cells can use juxtacrine signaling to exchange ROS with each other^57^. To use this route, however, the levels of ROS transported cannot be too high, as the transmitting or receiving cells could be damaged. Cells living within a biofilm, colony, and/or within different plant and animal tissues, are thought to use juxtacrine signalling to mediate cell-to-cell ROS and redox signaling^10,11^. Moreover, it was suggested that ROS can enhance the size of tunneling nanotubes or PDs and facilitate juxtacrine signalling^23,57^. Because we did not distinguish between paracrine and juxtacrine signalling in our study, we cannot rule out the possibility that the cells we imaged used one or the other, or both, for the process of cell-to-cell ROS signaling. In this respect it should be mentioned that our studies of the ROS wave in plants have shown that for cell-to-cell paracrine ROS signaling to occur, juxtacrine signalling must also occur, as key proteins localized to the PD must be present^23^. Paracrine and juxtacrine signalling might therefore be linked in other organisms and this aspect of cell-to-cell ROS signaling should be addressed in future studies.

Cancer cells use ROS to regulate their own proliferation and to communicate with cancer-associated macrophages and other host cells (*e.g*., ^58^). In addition, ROS plays a key role in cancer metastasis (*e.g*., ^59^). Our findings that cancer cells can generate and transmit long distance ROS signals (Figure 7), could suggest that a similar process may occur within the tumor microenvironment. This finding should be further studied as it may affect the development of new cancer fighting drugs. In the heart, ROS were implicated in ischemia reperfusion (IR) injury, both in the acute setting of myocardial infarction (MI), as well as in the chronic remodeling process following a MI (*e.g*., ^60–62^). Our findings that cardiomyocytes can use ROS for cell-to-cell communication (Figure 7; Movie S1), and that an active process of ROS signaling is triggered in isolated hearts upon localized wounding (Figure 8), could suggest that enhanced ROS production during IR could trigger different ROS signals that would augment or suppress the tissue damage caused during IR injury. This finding should also be further studied as it could lead to the development of new therapies that will limit tissue damage during IR.

Taken together, our findings suggest that cell-to-cell ROS signaling evolved before unicellular and multicellular organisms diverged (Figures 1–8)^19–24^. Because ROS are thought to have been present on Earth for billions of years and could be used to sense the cell’s environment through changes in the redox status of the cell^28,32,47,48^, it is possible that a cell-to-cell process that uses ROS as a signal evolved early in evolution. This mechanism could have communicated an environmental stress signal that altered the redox state of one cell (the initiating cell) through ROS secreted or transported by this cell to an acceptor cell (Figure 9). The changes in redox state caused by the perceived ROS signal in the acceptor cell would therefore have ‘mirrored’ the redox state of the initiating cell and information in the form of a ‘redox change’ transmitted from one cell to the other (Figure 9). The changes in redox state occurring in each of the two cells could cause ROS accumulation and activate gene expression through oxi-PTMs linking the gene expression and ROS responses of both cells (Figure 9). Since the redox state of other neighboring cells will also be altered in a similar manner during this process (like the redox changes that occur in cells along the path of the ROS wave in plants)^63^, and these cells will also start accumulating and secreting ROS, the entire colony, biofilm, or tissue would respond in unison to the stress perceived by the initiating cell (Figure 9), improving its chances of acclimation to the sensed stress, as well as overall survival. Such a mechanism could have coordinated the redox and ROS accumulation responses of a network of cells and give this cell population a better chance of survival in the early environment that was present on Earth; and may have been conserved in one form or another in different organisms to date (Figures 1–8)^19–24^. ROS such and H_2_O_2_ could have therefore served as one of the first ‘stress hormones’ on Earth; and to induce a response, the level of this stress hormone (*i.e*., H_2_O_2_) had to be increased (Figures 1–9). Our findings that an inhibitor of NADPH oxidases (DPI) can inhibit this process in different organisms (Figures 5–8), support the active nature of this process^19–21^. In addition, at least in *Chlamydomonas* we show that the cell-to-cell ROS signal is required for the expression of sHSP17.1 in cells that were not directly subjected to the stress (Figure 6), demonstrating that the systemic cell-to-cell ROS signal conveys important biological information between cells. In future studies it would be interesting to dissect this signaling process in different organisms and to determine whether it is dependent on paracrine, juxtacrine, or a combination of both signaling pathways (as in plants^20,21,23^). In addition, the nature of the ROS producing, transporting, and perceiving mechanisms need to be determined in different organisms. In plants these were found to be RBOHD/RBOHF, the aquaporin PIP2;1, and HPCA1, respectively^19–21,23,37^.

## Methods

### Organisms and stress treatments

The following organisms were studied in this research: *Pinus sylvestris* (TreeHelp, Buffalo, NY, USA); *Azolla filiculoides* (Carolina Biological Supply Company, Burlington, NC, USA), *Selaginella moellendorffii* (Putnam Hill nursery, Forest Hill, MD, USA), *Marchantia polymorpha* (Carolina Biological Supply Company, Burlington, NC, USA), *Anthoceros agrestis* (a kind gift from Prof. Fay-Wei Li, Cornell University), *Physcomitrium patens* (a kind gift from Prof. Elisabeth Haswell, Washington University at St. Louis and International Moss Stock Center, and Prof. Ralf Reski, University of Freiburg, Freiburg, Germany), *Chara vulgaris* (Carolina Biological Supply Company, Burlington, NC, USA), *Chlamydomonas reinhardtii* (UTEX Culture Collection of Algae, Austin, TX, USA), *Dictyostelium discoideum* (a kind gift from Prof. Thierry Soldati, University of Geneva, and Carolina Biological Supply Company, Burlington, NC, USA), and H9c2 and MDA-MB-231 (ATCC, Manassas, VA, USA). Organisms were grown according to stock resource instructions. Photosynthetic organisms were grown under 8 hrs/ 16 hrs light/dark regime. *P. sylvestris, S. moellendorffii and M. polymorpha* were grown in peat pellets (Jiffy-7, Jiffy International, Kristiansand, Norway)^20,21^. *A. filiculoides* was grown in 0.25 MS liquid media^64^; *A. agrestis* was grown on a solid modified Hatcher medium^65^; *P. patens* was grown on BCD solid media^66^; *Chara vulgaris* was grown in a Broyer and Barr liquid media^67^; *C. reinhardtii* cultures were grown on P49 solid media (UTEX Culture Collection of Algae, Austin, TX, USA); *D. discoideum* cultures were grown on SM agar plates in room temperature^68^. Mammalian cells were grown in 37°C 5% CO2 RPMI medium 1640 supplemented with 10% FCS, L-glutamine, and antibiotics (ThermoFisher, Waltham, MA)^69^. A day before the experiment the media was replaced with FluoroBrite DMEM (ThermoFisher, Waltham, MA) to reduce background fluorescence^70^. For organisms treated with local HL stress, following fumigation with H2DCFDA solution, illumination of 1700 μmol photons m^-2^ s^-1^ was applied to a local leaf/area for 2 min using a fiber optic (Schott, Southbridge, MA, USA), as described previously^20,23,27,37^. Heat injury, following treatment of H2DCFDA was applied for *D. discoideum* and mammalian cells using an open-flame-heated metal rod^22^ that touched the local injury area for 1 second^22^. Hydrogen peroxide was applied as a 1 μl drop of 10 mM diluted in the buffer of the medium in which the organism was cultured. Diphenyleneiodonium chloride (DPI; 50 μM) or water were applied as a strip of 0.05% agarose (applied with a pipette tip along the center of the plate; Figure 5A) for *P. patens, C. reinhardtii* and *D. discoideum*, or in the media for mammalian cells 5 or 10 min prior to the stress application^20^.

### Isolated heart studies

Mice used in this study were housed under pathogen-free conditions in animal care facilities and received humane care in compliance with the Guide for the Care and Use of Laboratory Animals (https://www.ncbi.nlm.nih.gov/books/NBK54050/)^71^. All experimental procedures were approved by the Institutional Animal Care and Use Committee of the University of Missouri (https://research.missouri.edu/acqa/acuc). C57BL/6J mice were obtained from Jackson Laboratory (https://www.jax.org/) at 8 weeks of age and used between the age of 9 and 10 weeks of age. Mice were placed under anesthesia using 4% Isoflurane, sacrificed, and the hearts were immediately collected^71^. The hearts were rinsed in FluoroBrite DMEM (ThermoFisher, Waltham, MA) to remove excess blood. They were then incubated in Fluro Bright DMEM media containing 10% Fetal Bovine Serum, 15 mM Hepes, 50 μM H2DCFDA (Millipore-Sigma, St. Louis, MO, USA), and Penicillin-Streptomycin 10,000 Units/ml (Gibco; ThermoFisher, Waltham, MA) for 2 hrs with gentle rocking in a CO_2_ incubator (Binder C170; ThermoFisher, Waltham, MA). For DPI treatments, DPI (50 μM final concentration) was added to the incubation media 30 min prior to imaging. Isolated hearts were then wounded or unwounded with an open-flame-heated rod^22^ and immediately imaged as described below.

### ROS imaging

All organisms, outside of mammalian cells and isolated hearts, were fumigated with 50 μM H2DCFDA (Millipore-Sigma, St. Louis, MO, USA) in 0.05 M Phosphate buffer pH 7.4 with 0.01% Silwet L-77. Fumigation was carried out for 30 min using nebulizers (Punasi Direct, Hong Kong, China)^20–24,37,38^. For mammalian cells and isolated hearts 10 μM H2DCFDA (final concentration) was applied to the medium for 30 min (cell cultures), or 2 hrs (isolated hearts). Following H2DCFDA fumigation/incubation, local stress/wounding was applied, and organisms/isolated hearts were imaged for live ROS accumulation using the IVIS Lumina S5 platform^20–24,37,38,72^. Acquired images were then analyzed using the math function in Living Image software (PerkinElmer, Waltham, MA, USA) ^20–24,37,38^. Empty agar plates were also fumigated with 2’,7’-dichlorodihydrofluorescein (DCF; OxyBurst; 50 μM)^19^, treated with H_2_O_2_ and imaged as described above.

### Protein blot analysis

Local and systemic cells of *C. reinhardtii* from treated or untreated plates, with or without DPI, 2 min or 30 min after a 2 min HL treatment were collected and denatured by boiling in 4 x Laemmli buffer. The samples were spun down and the supernatant was loaded on SDS-PAGE and blotted to PVDF membrane. Polyclonal anti-sHSP17.1 antibodies from rabbit were used to identify the proteins on the membrane as described previously^24,25^.

### Statistical analysis

All experiments were repeated at least three times with at least 3 biological repeats. Box plots graphs are presented with mean as X ± SE; median is line in the box and box borders are 25th and 75th percentiles; whiskers are the 1.5 interquartile range. Paired Student t-test was conducted using Microsoft Excel (*P < 0.05; **P < 0.01; ***P < 0.001).

## Acknowledgements

We thank Professor Brent Mishler for discussions regarding which organisms to target for our evolutionary analysis. We also thank Professor Thierry Soldati for gift of *Dictyostelium discoideum*, Professor Elisabeth Haswell and Professor Ralf Reski for gift of *Physcomitrium patens*, and Professor Fay-Wei Li for gift of *Anthoceros agrestis*.

## Figure legends

**Table S1.**
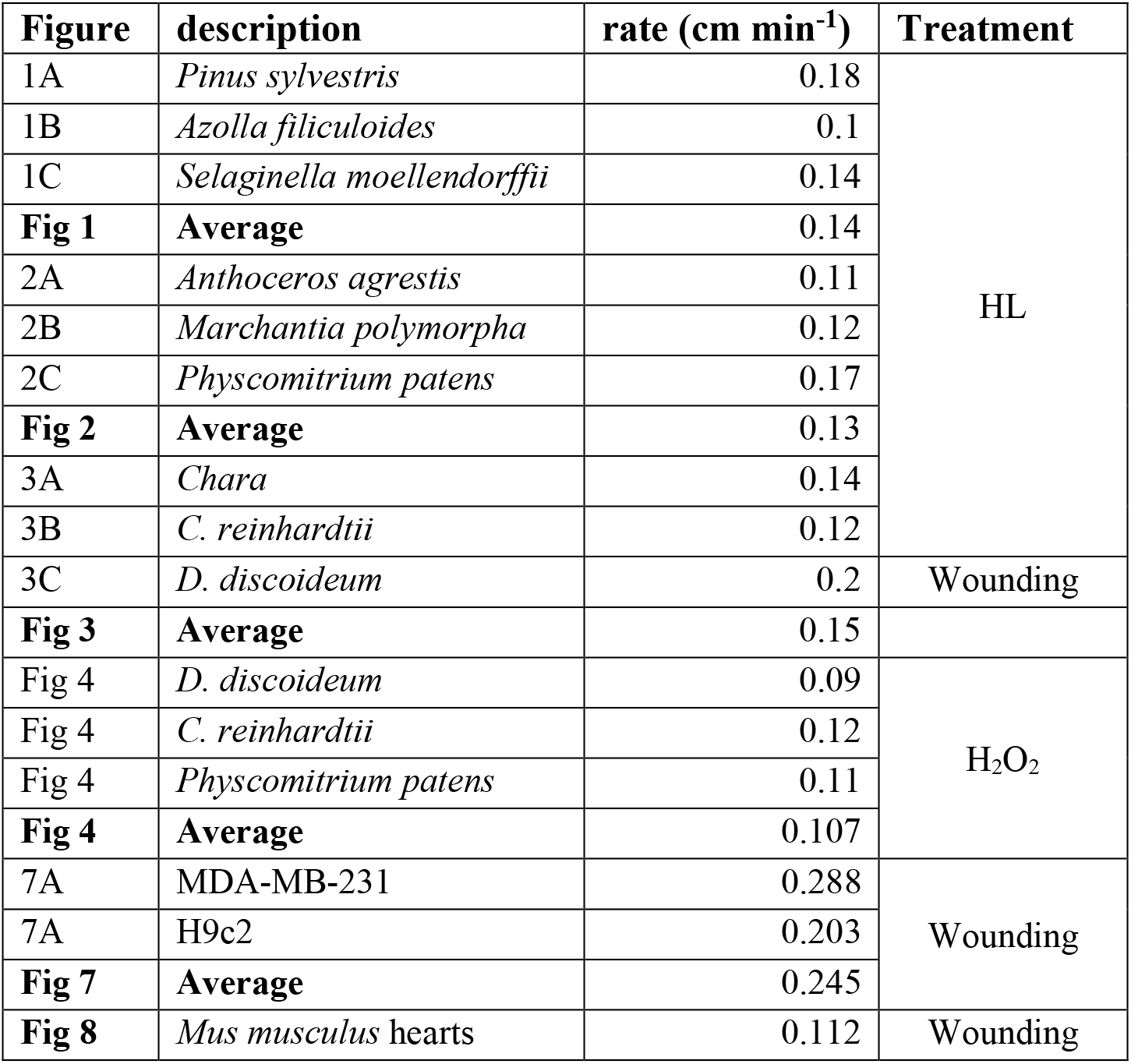
Rates of the ROS wave in the different experiments.

**Figure S1.**
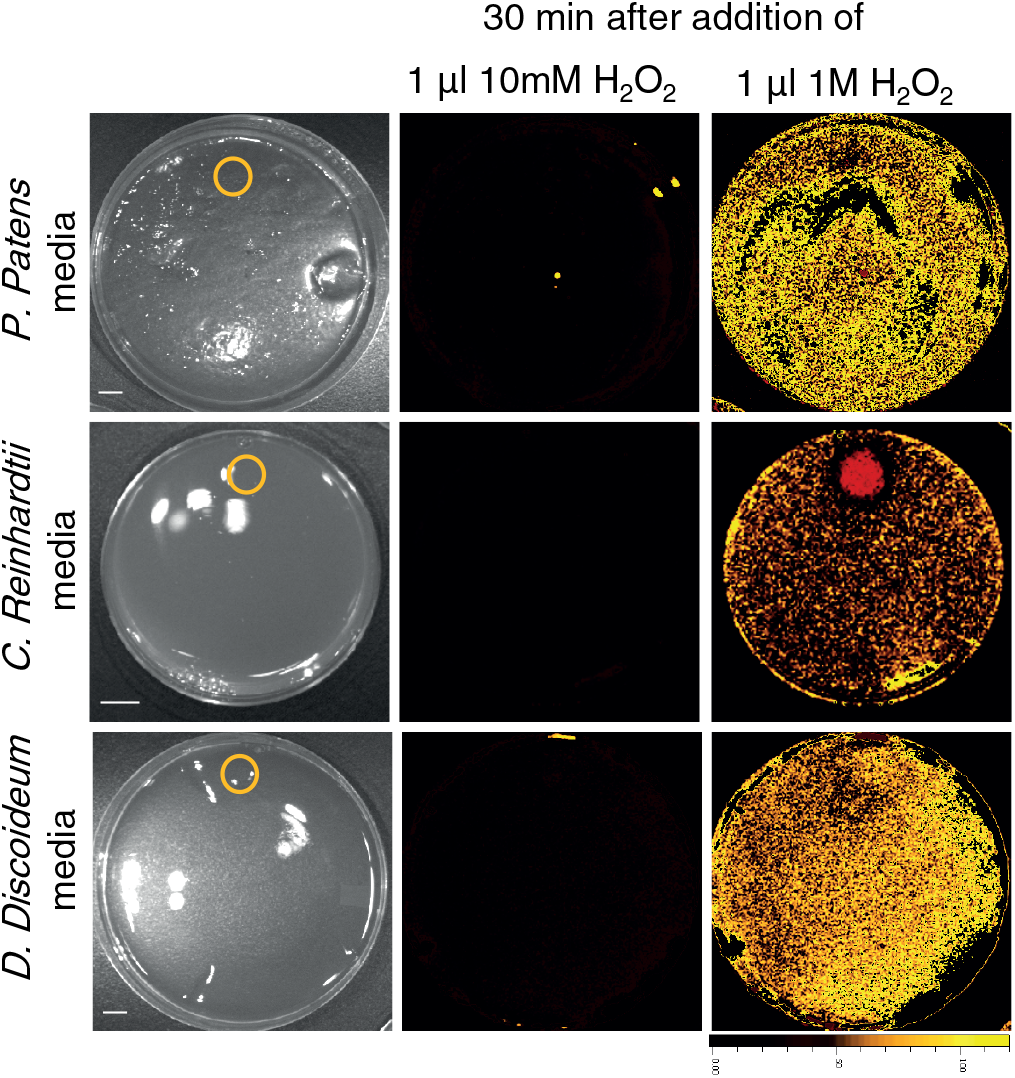
Controls for the detection of H_2_O_2_ on agar plates (in support of Figure 4). Agar plates containing the different growth media used in Figure 4, but without any organisms growing on them, were fumigated with OxyBurst, which detects ROS in the absence of cells. A 1 μL drop of 10 mM or 1 M H_2_O_2_ was then applied to the plate (indicated with a circle) and whole plate ROS accumulation was imaged for 30 min. Scale bar, 1 cm.

**Figure S2.**
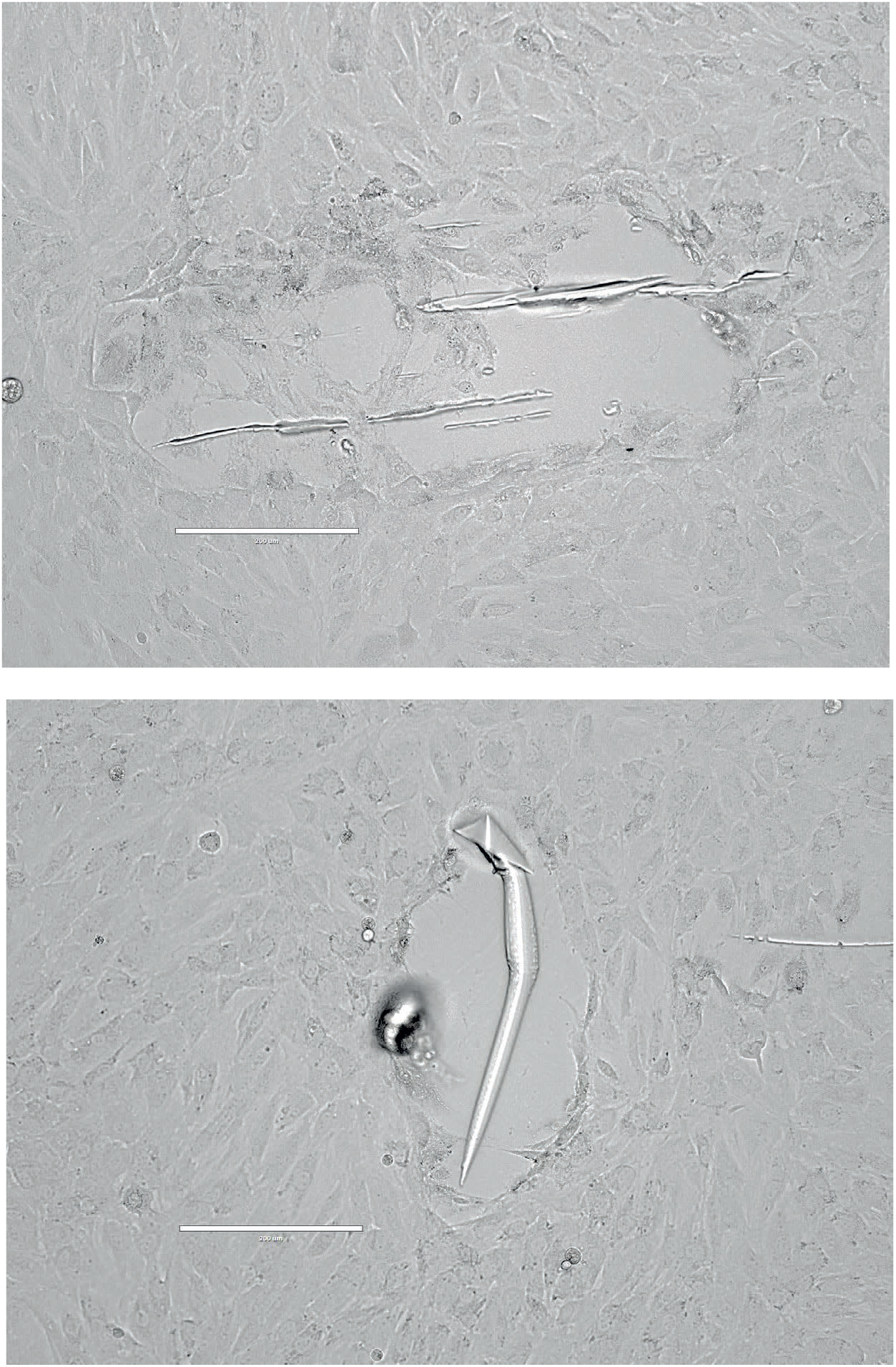
Representative images of injury inflicted on the H9c2 cell monolayer by a heat-treated metal rod (in support of Figure 7a). Plates with a monolayer of H9c2 cells, used for the experiments shown in Figure 7a were imaged with an Agilent BioTek Lionheart FX automated microscope and the different images collected were assembled to reveal the size of injury inflicted on the monolayer of cells. Scale bar, 200 μM.

**Figure S3.**
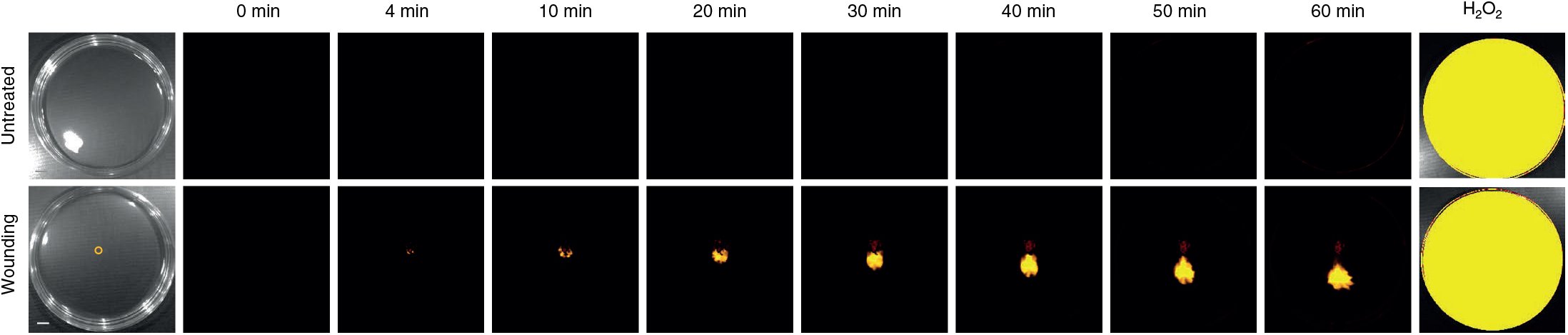
Controls for the ability of H9c2 cells that do not show ROS accumulation following the heated rod injury to accumulate ROS (in support of Figure 7). Monolayers of rat cardiomyocytes (H9c2), cells grown in culture were treated or untreated with a heated metal rod (indicated with a circle) to induce injury (average injury diameter is 120 μm) and ROS accumulation was imaged using H2DCFDA for 60 min in the entire plate. At the end of the 60 min, H_2_O_2_ at a final concentration of 1 mM was added to the plates and the plates were imaged for an additional 10 min. Representative time-lapse images of whole plate ROS accumulation in treated and untreated plates are shown. Scale bar, 1 cm. Abbreviations: H_2_DCFDA, 2’,7’-dichlorodihydrofluorescein diacetate; ROS, reactive oxygen species.

**Figure S4.**
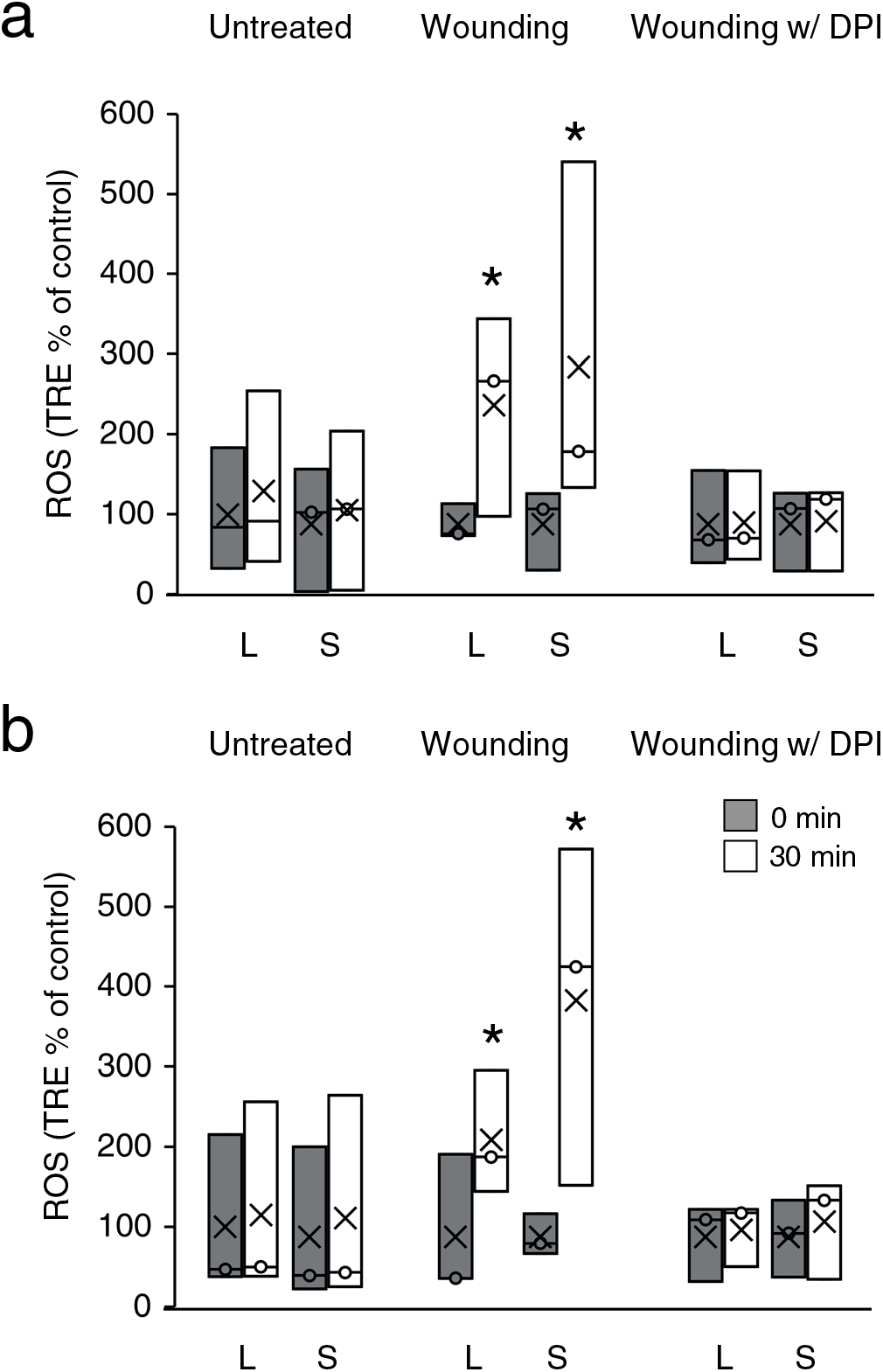
Wound-induced rapid systemic ROS accumulation in hearts from female and male mice. Mice hearts, surgically removed from 9-10-week-old male and female C57BL/6J mice, were untreated, or treated with a heated metal rod (to induce injury), in the presence or absence of DPI (50 μM), and ROS accumulation was measured using H2DCFDA as described in Figure 8. Bar graphs of combined data from all female **(a)** and male **(b)** hearts used for the analysis at the 0- and 30-min time points are shown (in support of Figure 8). All experiments were repeated at least 3 times with 3 female and 3 male mice per experiment. Data is presented as box plot graphs; X is mean ± S.E., N=30, **P < 0.01, ***P < 0.001, Student t-test. Scale bar, 1 cm. Female- and male-specific ROS responses are shown in Figure S4. Abbreviations: DPI, diphenyleneiodonium; H2DCFDA, 2’,7’-dichlorodihydrofluorescein diacetate; L, local; ROS, reactive oxygen species; S, systemic; TRE, total radiant efficiency; w/, with.

**Movie S1.** A movie showing the ROS signal generated in a monolayer of H9c2 cells following heat injury (in support of Figure 7a). Monolayers of rat cardiomyocytes (H9c2) cells grown in culture were treated with a heated metal rod (indicated with a circle) to induce injury (average injury diameter is 120 μm) and ROS accumulation was imaged using H_2_DCFDA in the entire plate. Abbreviations: H_2_DCFDA, 2’,7’-dichlorodihydrofluorescein diacetate.

**Movie S2.** A movie showing the ROS signal generated in an isolated mice heart following heat injury (in support of Figure 8a). Isolated hearts from mice were treated with a heated metal rod (indicated with a circle) to induce injury and ROS accumulation was imaged using H_2_DCFDA in the entire plate. Abbreviations: H_2_DCFDA, 2’,7’-dichlorodihydrofluorescein diacetate.

